# Targeted genome fragmentation with CRISPR/Cas9 improves hybridization capture, reduces PCR bias, and enables efficient high-accuracy sequencing of small targets

**DOI:** 10.1101/207027

**Authors:** Daniela Nachmanson, Shenyi Lian, Elizabeth K. Schmidt, Michael J. Hipp, Kathryn T. Baker, Yuezheng Zhang, Maria Tretiakova, Kaitlyn Loubet-Senear, Brendan F. Kohrn, Jesse J. Salk, Scott R. Kennedy, Rosa Ana Risques

## Abstract

Current next-generation sequencing techniques suffer from inefficient target enrichment and frequent errors. To address these issues, we have developed a targeted genome fragmentation approach based on CRISPR/Cas9 digestion. By designing all fragments to similar lengths, regions of interest can be size-selected prior to library preparation, increasing hybridization capture efficiency. Additionally, homogenous length fragments reduce PCR bias and maximize read usability. We combine this novel target enrichment approach with ultra-accurate Duplex Sequencing. The result, termed CRISPR-DS, is a robust targeted sequencing technique that overcomes the inherent challenges of small target enrichment and enables the detection of ultra-low frequency mutations with small DNA inputs.

## BACKGROUND

In the past decade, NGS has revolutionized the fields of biology and medicine. However, standard NGS suffers from two major problems that negatively impact multiple applications: the limited efficiency of the current target selection methods and the high error rate of the sequencing process. Targeted genome enrichment is essential to many applications that do not require whole genome sequencing and it is performed either by PCR or by hybridization capture. PCR is simple and efficient but does not scale well and suffers from biases that result in uneven coverage and false mutation calls [1, 2]. Hybridization capture improves coverage uniformity and mutation call accuracy but has low recovery, especially when the target region is small, which leads to the requirement of larger amounts of DNA [2]. An additional complication is that DNA is typically fragmented by sonication which introduces DNA damage resulting in sequencing errors [3]. Moreover, the heterogeneous fragment sizes generated by sonication are subject to PCR bias and contribute to uneven coverage. An alternative option to sonication is enzymatic fragmentation. This method resolves some issues but introduces different artifacts that also result in sequencing errors [4]. Thus, at the library preparation step, both methods of target selection suffer important limitations that lead to non-optimal sequencing outcomes, including uneven coverage, introduction of false mutations, and low recovery.

The second major problem of NGS is the high error rate inherent to the sequencing process. Illumina offers the most accurate sequencing platform with an estimated error rate of 10^−3^ [5]. This error rate, however, translates into millions of false calls in each sequencing run and precludes the detection of low frequency mutations (Additional file 1: Figure S1), which is critical for applications such as forensics, metagenomics, and oncology [6]. While the accuracy of NGS can be improved by repair of the DNA prior to sequencing [7, 8] and by computational error correction [9, 10], these strategies do not remove all potential artifacts. An alternative approach employs unique molecular identifiers, also called molecular barcodes or molecular tags, to identify the reads derived from an original DNA molecule and use their redundant information to create a consensus sequence [11]. The unique random shear points generated at sonication can be used as “endogeneous barcodes”, but only when sequencing depth is low (~10x) to avoid overlapping of sharing points between independent DNA molecules [12]. To enable higher sequencing depth, exogenous barcodes are necessary. Exogenous barcodes are random DNA sequences attached to the original DNA molecules before or during PCR. Single-stranded molecular barcodes produce a consensus with the reads derived from one DNA strand [11], whereas double-stranded molecular barcodes introduce an additional level of correction by allowing the comparison of independent consensus sequences derived from the two complementary strands of the original DNA molecule [13]. This additional level of correction is essential for removing polymerase errors occurring in the first round of PCR and subsequently propagated to all reads derived from a given DNA strand [7]. Polymerase errors caused by DNA damage are one of the most pervasive problems of NGS [8] but can be successfully addressed with double-strand barcodes given the extremely low probability that the same error occurs in the same position on both strands of DNA. Duplex Sequencing (DS), the method that pioneered double-strand molecular barcodes [13, 14], has an estimated error rate <10^−7^, four orders of magnitude less than single-strand molecular barcode methods. This level of accuracy allows for very sensitive ultra-deep sequencing (Additional file 1: Figure S1) and has been employed in a variety of applications including the detection of very low frequency somatic mutations in cancer and aging [15-18].

DS successfully addresses the problem of sequencing errors, but it suffers from the limitations of hybridization capture, which is required to perform target selection while preserving the strand recognition of molecular barcodes. As described above, hybridization capture is highly inefficient when selecting small target sizes [19]. It is estimated that for targets <50Kb only 5-10% of reads are on-target after hybridization capture [20]. In DS, as well as in other panel-based sequencing approaches, the region of interest is usually small as a cost-effective trade-off for higher sequencing depth. In this situation, a successful approach for target enrichment is to perform two consecutive rounds of capture [20]. However, this approach results in a time consuming, costly, and inefficient protocol that requires large amounts of DNA [14]. For example, in DS at least 1μg of DNA was historically needed to produce depths >3,000x [17], which is prohibitive in many applications that rely on small samples.

Here we present CRISPR-DS, a new method that addresses the two main problems of NGS: limited efficiency of target selection and high error rate. Target selection is facilitated by an enrichment of the regions of interest using the CRISPR/Cas9 system. *In vitro* digestion with CRISPR/Cas9 has been proven to be a useful tool for multiplexed excision of large megabase fragments and repetitive sequence regions for PCR-free NGS [21, 22]. We reasoned that targeted *in vitro* CRISPR/Cas9 digestion could be used to excise similar length fragments covering the area of interest, which could then be enriched by size selection prior to library preparation. We designed this method to enable target enrichment while simultaneously eliminating sonication-related errors and biases arising from random genome fragmentation. In addition, by pairing this method with double-strand molecular barcoding, we aimed to produce a method that preserves the sequencing accuracy of DS, while increasing the recovery rate, enabling low DNA input and a simplified protocol for translational applications.

## RESULTS

### Design of CRISPR-DS based on CRISPR/Cas9 target fragmentation and double strand molecular barcodes

CRISPR-DS is based on *in vitro* CRISPR/Cas9 excision of target sequences to generate DNA molecules of uniform length which are then enriched by size selection. The versatility, specificity, and multiplexing capabilities of the CRISPR/Cas9 system enable its application for the excision of any target region of interest by simply designing guide RNAs (gRNA) to the desired cutting points. As a proof of principle, we developed the method for sequencing the exons of *TP53*. Further, in order to achieve high recovery as well as high sequencing accuracy, we combined it with DS. The main steps of the protocol are illustrated in Figure 1. First, target regions are excised from genomic DNA by multiplexed *in vitro* CRISPR/Cas9 digestion (Fig. 1a), followed by enrichment of the excised fragments by size-selection using SPRI beads (Fig. 1b). The selected fragments are then coupled with the double-strand molecular barcodes used in DS (Fig. 1c) [14]. These fragments are then amplified and captured with biotinylated hybridization probes as previously described for DS [14], with the exception that only one round of hybridization capture is required due to the prior enrichment of target fragments (see below). Finally, the library is sequenced and the resulting reads are analyzed to perform error correction based on the consensus sequences of both strands of each DNA molecule (Fig. 1d) [14]. Due to the requirement of only one round of hybridization capture, the workflow of CRISPR-DS is almost one day shorter than standard-DS (Fig. 2, Additional file 1: Figure S2), enabling a more cost-efficient and applicable method.

**Figure 1.**
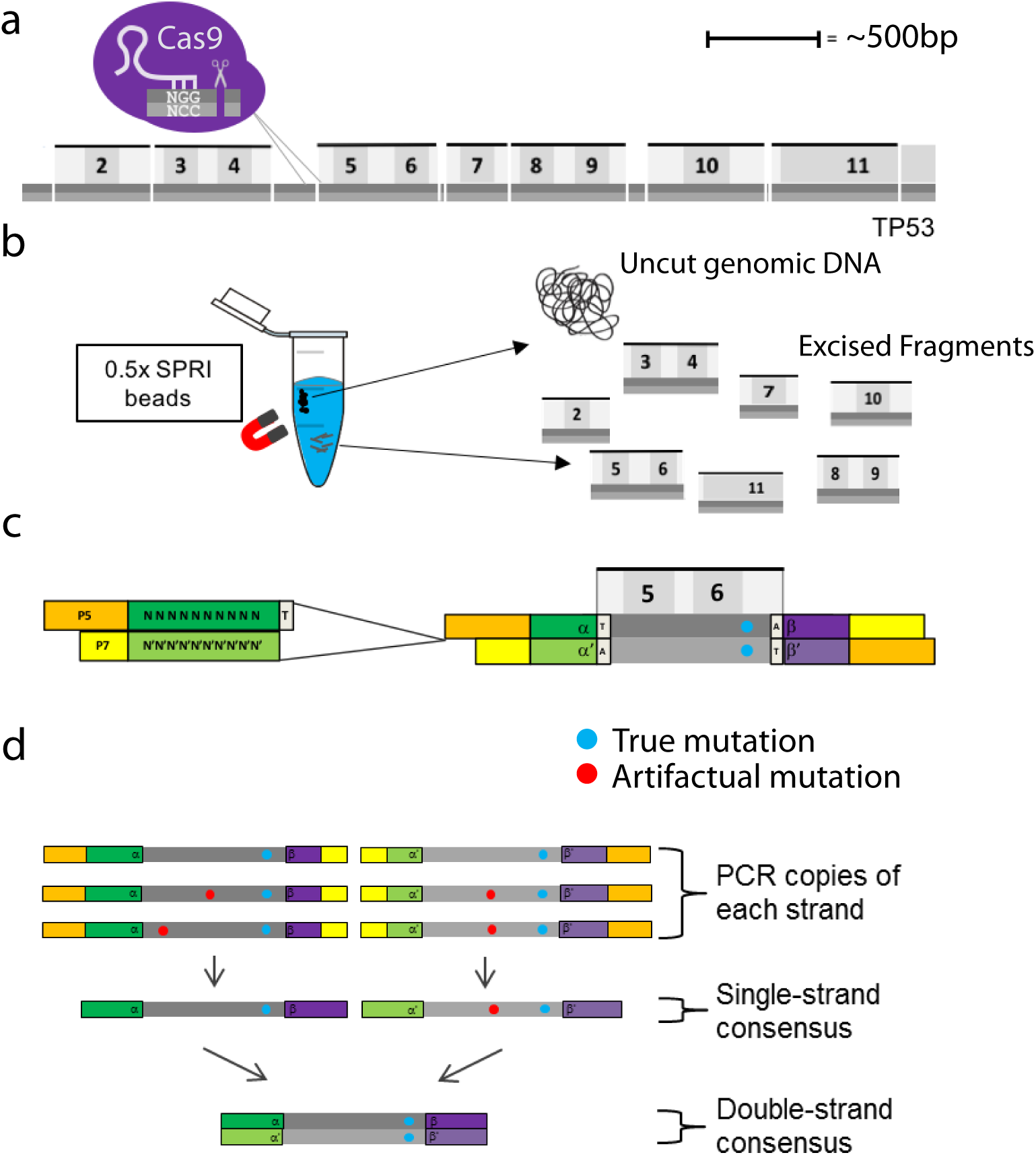
Schematic representation of key aspects of CRISPR-DS. (a) CRISPR/Cas9 digestion of *TP53*. Seven fragments containing all *TP53* coding exons were excised via targeted cutting using gRNAs. Dark grey represents reference strand and light grey represents the anti-reference strand. (b) Size selection using 0.5x SPRI beads. Uncut, genomic DNA binds to the beads and allows the recovery of the homogenously sized excised fragments in solution. (c) Double-stranded DNA molecule fragmented and ligated with DS-adapters. Adapters are double-stranded and contain 10-bp of random, complementary nucleotides and a 3’-dT overhang. (d) Error correction by DS. Reads derived from the same strand of DNA are compared to form a Single-Strand Consensus Sequence (SSCS). Then both strands of the same original DNA molecule are compared with one another to create a Double-Strand Consensus Sequence (DCS). Only mutations found in both SSCS reads are counted as true mutations in DCS reads.

**Figure 2.**
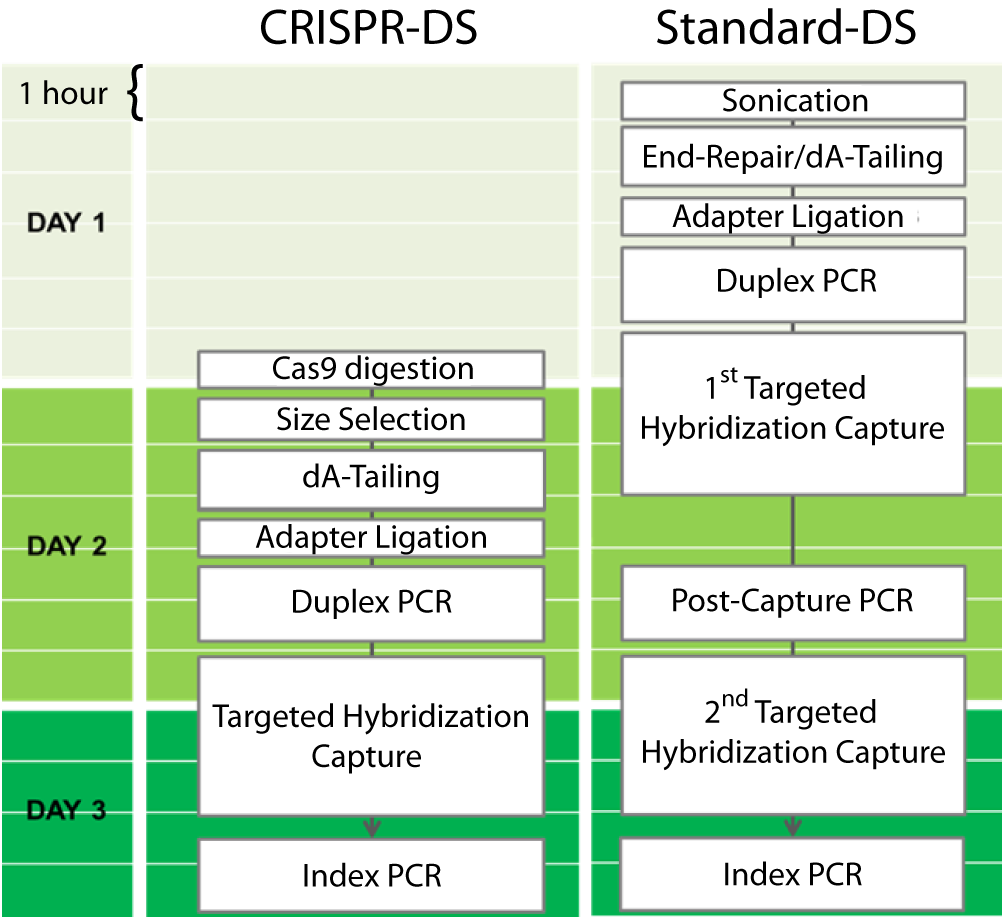
Comparison of library preparation protocols for standard-DS vs. CRISPR-DS. The primary differences between the CRISPR-DS and standard-DS library preparation are the fragmentation techniques and the number of hybridization capture steps. Instead of fragmentation by sonication as performed in standard-DS, CRISPR-DS relies on an *in vitro* excision of target regions by CRISPR/Cas9 followed by size selection for the excised fragments. The size selection eliminates the need for a second round of hybridization capture which is required for sufficient target enrichment in the standard-DS protocol. CRISPR-DS reduces the workflow by nearly a day. Colored boxes represent 1h of time.

### CRISPR/Cas9 cut fragments can be designed to be of homogenous length, reducing PCR bias and producing uniform coverage

Typically, genome fragmentation is performed with sonication, which generates randomly sized fragments that have different amplification efficiencies [23]. Short fragments are preferentially amplified, resulting in uneven coverage of the regions of interest and decreased recovery. In DS, amplification bias introduces an additional problem because short fragments produce an excess of PCR copies that do not further aid error reduction. To produce a consensus, only three PCR copies of the same molecule are required. Additional copies waste resources because they produce sequencing reads but do not generate additional data. By using CRISPR/Cas9, gRNA can be designed such that restriction with Cas9 produces fragments of predefined, homogeneous size. We reasoned that these fragments would eliminate PCR bias, leading to homogeneous sequencing coverage and minimizing wasted reads that are PCR copies of the same original molecule.

To test this approach, we designed gRNAs to specifically excise the coding regions and their flanking intronic sequence of *TP53* (Fig. 1a). Fragment length was designed to be ~500bp in order to maximize read space of an Illumina MiSeq v3 600 cycle kit while allowing for sequencing of the molecular barcode (10 bp) and 3’-end clipping of 30bp to remove low-quality bases produced in the later sequencing cycles. gRNAs were selected based on the highest specificity score that produced appropriate fragment length (Additional file 2: Table S1, Additional file 3: Data S1) [24]. The fragment comprising exon 7 was designed shorter than the rest (336 bp) to avoid a homopolymeric run of T’s in the flanking intronic region which induced poor base quality in reads that span this region (Additional file 1: Figure S3).

We performed a side-by-side comparison of library performance (Fig. 3a-c) and sequencing coverage (Fig. 3d) of a sample DNA processed with CRISPR-DS vs standard-DS (see Material and Methods). Standard-DS for *TP53* had been previously performed using sonication and published protocols [14, 17]. Visualization of the resulting sequencing library by gel electrophoresis showed that CRISPR restriction produced distinct bands/peaks (Fig. 3a-b) corresponding to the predesigned size of target fragments as opposed to the diffuse “smear” characteristic of libraries prepared by sonication. The discrete peaks allow confirmation of correct library preparation and target enrichment, preventing the sequencing of suboptimal libraries. Sequencing and mapping of the libraries demonstrated that targeted Cas9 restriction results in well-defined DNA fragments corresponding to the expected size (Fig. 3d). Importantly, these fragments exhibited extremely uniform sequencing depth. In contrast, sonicated DNA fragments resulted in significant variability in depth across target regions. Because DS reads correspond to individual DNA molecules, the uniform depth achieved by CRISPR-DS indicates a homogenous representation of the original genomic DNA in the final sequencing output, confirming the proper excision of all fragments.

**Figure 3.**
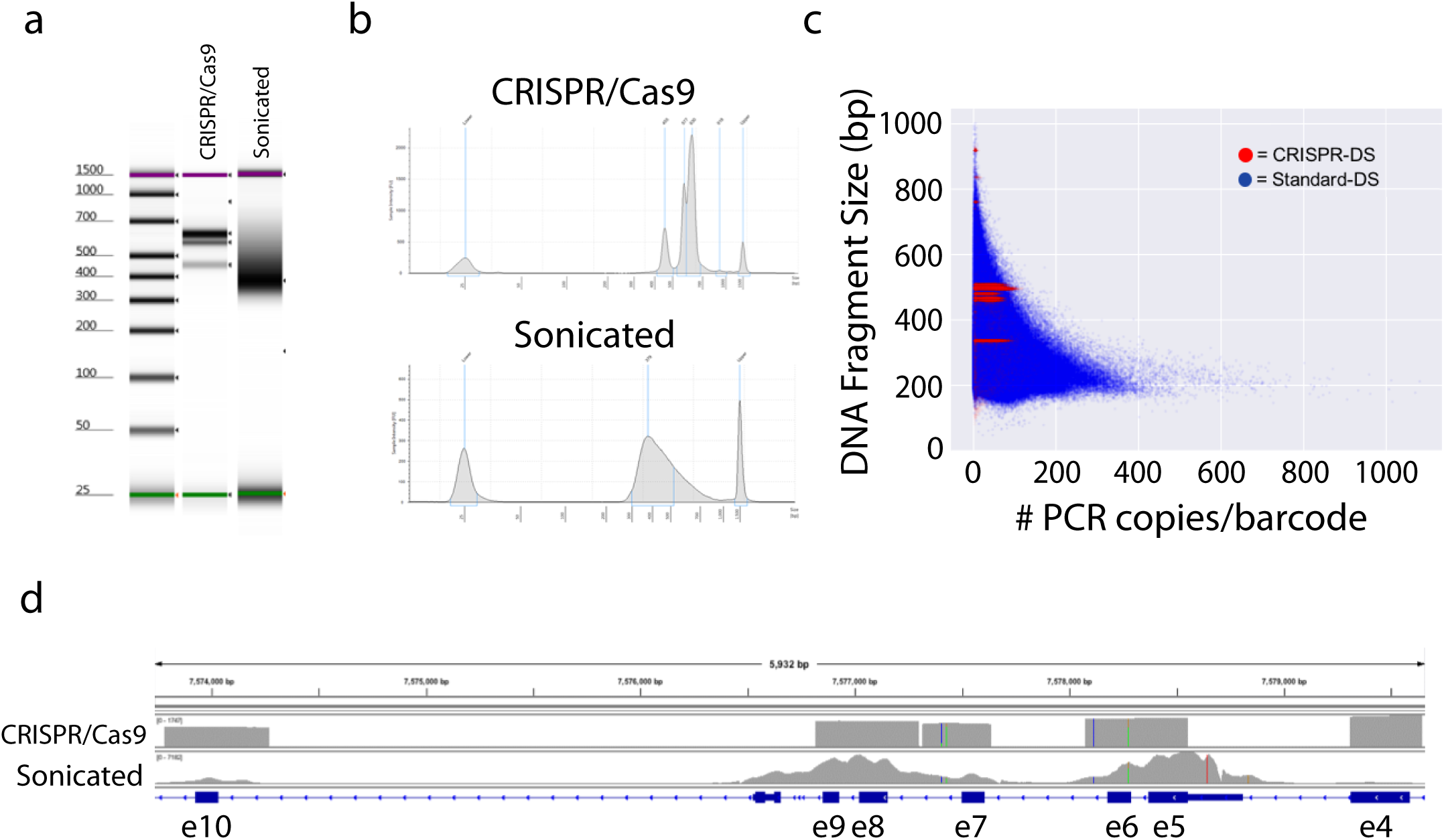
Visualization of sequencing libraries and data prepared with CRISPR-DS and standard-DS. (a) TapeStation gels show distinct bands for CRISPR-DS as opposed to a smear for standard-DS. The size of bands corresponds to the CRISPR/Cas9 cut fragments with adapters. (b) CRISPR-DS electropherograms allow visualization and quantification of peaks for quality control of the library prior to sequencing. Standard-DS electropherograms show a diffuse peak that harbors no information about the specificity of the library. (c) Dots represent original barcoded DNA molecules. Each DNA molecule has multiple copies generated at PCR (x-axis). In CRISPR-DS, all DNA molecules (red dots) have preset sizes (y-axis) and generate similar number of PCR copies. In standard-DS, sonication shears DNA into variable fragment lengths (blue dots). Smaller fragments amplify better and generate an excess of copies that waste sequencing resources. (d) Integrative Genomics Viewer of *TP53* coverage with DCS reads generated by CRISPS-DS and standard-DS. CRISPR-DS shows distinct boundaries that correspond to the CRISPR/Cas9 cutting points and an even distribution of depth across positions, both within a fragment and between fragments. Standard-DS shows the typical ‘peak’ pattern generated by random shearing of fragments and hybridization capture, which leads to variable coverage.

The ability to uniformly control the DNA insert size should not only provide homogenous depth, but should also produce a more uniform number of copies of each molecule, minimizing the waste of unnecessary reads to produce a consensus sequence. To test this possibility, we counted the number of PCR copies for each molecular barcode and plotted it as a function of the DNA fragment size (Fig. 3c). Sonicated DNA exhibited a strongly negative association between DNA fragment size and the number of PCR copies, as expected due to the fact that small DNA fragments are preferentially amplified (Fig. 3c, *blue*). In contrast, targeted fragmentation produced a consistent number of PCR copies for all fragments, including the smaller exon 7 fragment (Fig. 3c, *red*).

### CRISPR/Cas9 cut fragments can be designed to be of optimal length to maximize read usage

An additional disadvantage of the variable fragment size produced by sonication is inefficient read usage: fragments that are too short generate overlapping reads that waste sequencing space, whereas fragments that are too long get sequenced on the ends, leaving captured but un-sequenced DNA in the middle (Fig. 4a). The programmable nature of Cas9 can be leveraged to reduce the amount of data “lost” by generating optimal length fragments tailored to the preferred number of sequencing cycles. To illustrate the improvement in read usage, we quantified the amount of deviation from the optimal fragment size (defined as the total number of sequencing cycles minus the total length of the molecular barcodes and 3’-end clipping) of seven samples independently processed with sonication and targeted fragmentation. Sonication produced significant variability in the amount of deviation from the optimal fragment size with a large fraction of fragments being twice the optimal size for one of the samples (Fig. 4b,c; Additional file 1: Figure S4). Indeed, only 9.1±4.2% of reads had inserts that were within 10% deviation from the optimal fragment length. Even samples with more stringent size selection had only ~61% of reads within the 10%-deviation window (Fig. 4c; Additional file 1: Figure S4). In contrast, the same samples fragmented with Cas9 had 71.0±3.2% of reads within the same window range, with the vast majority of the reads outside the window being due the purposefully shorter Exon 7 fragment (Fig. 4b,c; Additional file 1: Figure S3, S4). Exclusion of exon 7 from this analysis improved the percent of reads within the 10%-deviation window to 94.3±2.1%. These data indicate that targeted fragmentation can tightly control the fragment size to optimize read usage, thereby increasing the efficiency of sequencing.

**Figure 4.**
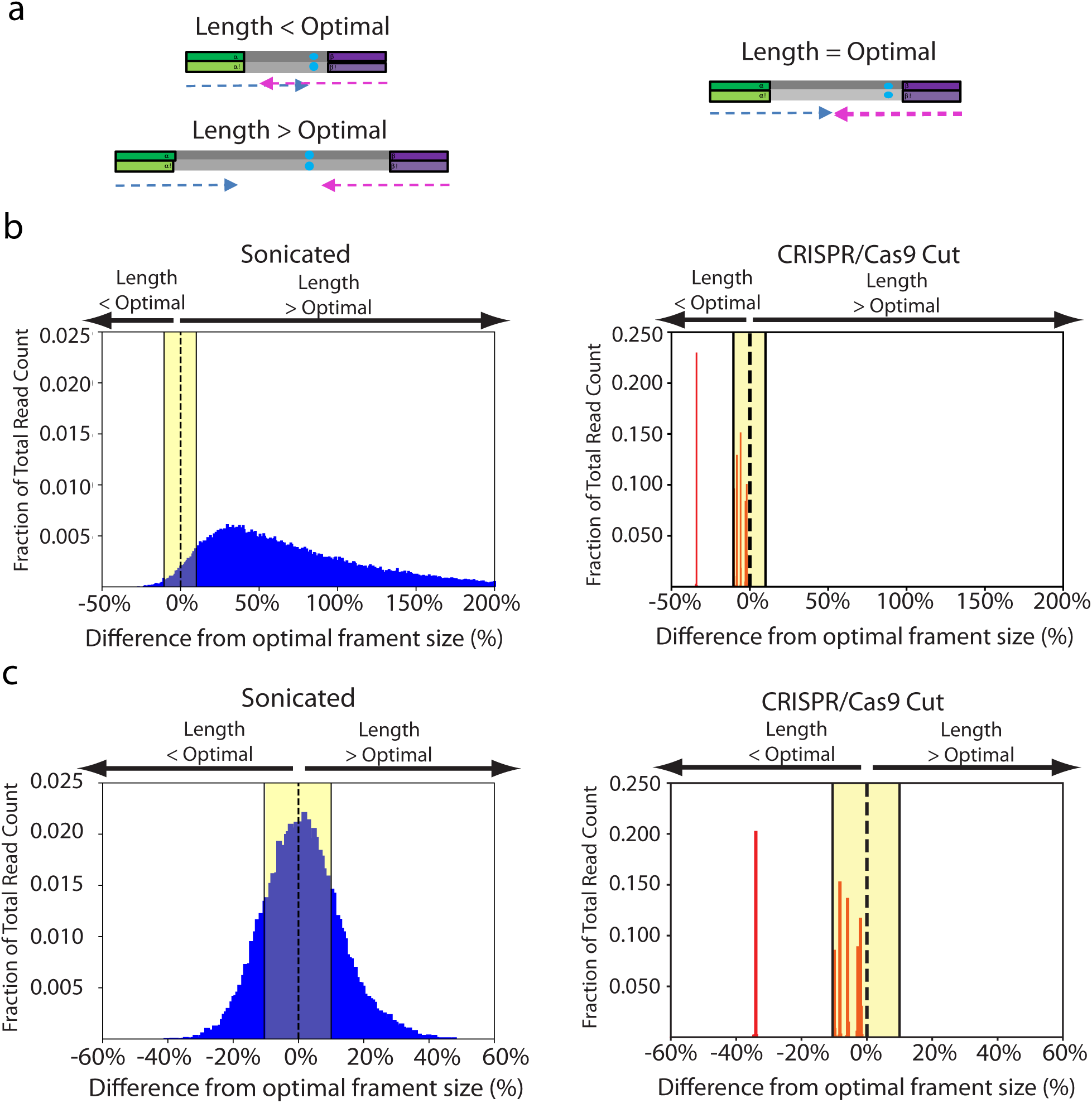
CRISPR/Cas9 fragmentation produces optimal fragment lengths. (a) Sonication produces fragments that are either too short or too long, corresponding to redundant or lost information, respectively. CRISPR-DS produces optimally sized fragments which are perfectly covered by the sequencing reads. (b-c) Comparison of histograms of the insert sizes of two samples prepared with standard-DS (*blue*, left panels), which uses sonication for fragmentation, and CRISPR-DS (*red*, right panels), which uses CRISPR/Cas9 digestion for fragmentation, The x-axis represents the percent difference from the optimally sized fragment, e.g. fragment size that matches the sequencing read length after adjustments for molecular barcodes and clipping. Yellow shading highlights range of fragment sizes which are within 10% difference from optimal size.

### CRISPR/Cas9 fragmentation enables target enrichment by size selection, eliminates one round of hybridization capture, and increases sequencing yield

While performing two rounds of capture substantially increases the number of on-target reads for standard-DS and other small target applications, the process is time consuming, expensive, and requires additional PCR steps that introduce further bias [20]. We hypothesized that target enrichment via size selection of CRISPR/Cas9 digested fragments would sufficiently enrich for on-target DNA fragments and eliminate the need for a second capture. To test this hypothesis, we performed CRISPR/Cas9 digestion of targeted *TP53* exons (Fig. 1a) on a range of DNA input amounts (10-250ng) followed by SPRI size selection to remove undigested high molecular weight DNA fragments (> 1kb in size). The selected DNA fragments were ligated to DS adapters, PCR amplified, and sequenced (see Material and Methods). No hybridization capture or any other type of target enrichment was performed. Mapping of raw reads revealed between 0.2% to 5% reads on-target, corresponding to ~2,000x to 50,000x fold enrichment given the fact that our target region only amounted for 0.000101% of the human genome (Table 1). This level of enrichment matches or exceeds what is typically achieved with solution based hybridization for small target size [19, 20]. Notably, lower DNA inputs showed the highest enrichment, potentially reflecting more efficient digestion or improved removal of off-target, high molecular weight DNA fragments when they are in lower abundance. These results suggested that a simple size selection step can be used in lieu of a targeted hybridization enrichment step.

**Table 1:** Target enrichment due to size selection

| Sample | DNA Input<br>(ng) | Reads On<br>Target (%) | Fold<br>Enrichment |
| --- | --- | --- | --- |
| B9 | 25 | 0.76% | 7,527 |
|  | 200 | 0.25% | 2,452 |
|  | 250 | 0.21% | 2,037 |
| PF1 | 10 | 2.85% | 28,139 |
|  | 25 | 1.99% | 19,583 |
|  | 100 | 0.68% | 6,667 |
|  | 250 | 0.70% | 6,878 |
| PF5 | 10 | 5.05% | 49,794 |
|  | 25 | 0.96% | 9,456 |
|  | 100 | 0.34% | 3,321 |
|  | 250 | 0.22% | 2,217 |

To test this possibility, we performed a side by side comparison of standard-DS [14] with one and two rounds of hybridization capture vs. CRISPR-DS with only one round of hybridization capture. Three input amounts of the same control DNA extracted from normal human bladder tissue were sequenced in parallel for each of the methods. CRISPR-DS with one round of capture achieved >90% raw reads on-target (e.g. covering *TP53*) (Fig. 5a), a significant improvement over standard-DS which only achieved ~5% raw reads on-target with a single capture, consistent with prior work [20]. In an independent experiment, we tested the reproducibility of this result with three different DNA samples that were sequenced with CRISPR-DS using one and two rounds of capture (Additional file 1: Figure S5). Confirming the prior result, the three samples produced >90% raw reads on target using only one round of capture. The second round of capture only minimally increased raw reads on-target and is unnecessary.

**Figure 5.**
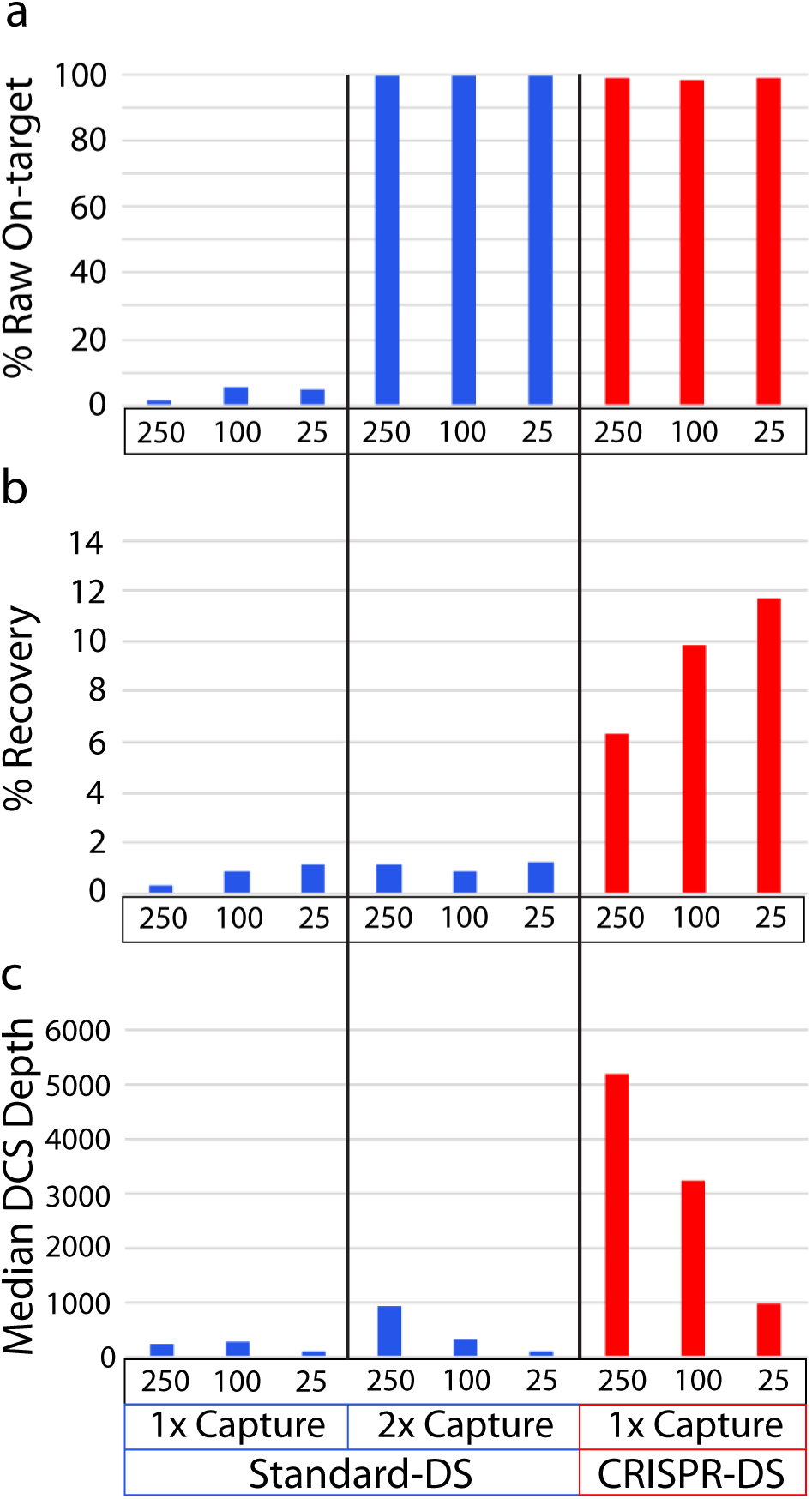
Technical comparison of 250ng, 100ng and 25ng of DNA sequenced with both standard-DS and CRISPR-DS. Measurements were obtained by sequencing samples prepared with standard-DS (*blue*) using one and two rounds of hybridization capture and CRISPR-DS (*red*) with only one round of hybridization capture. (a) The percentage of raw sequencing reads on-target (covering *TP53*) was comparable between Standard-DS with two rounds of capture and CRISPR-DS with one round of capture, demonstrating the target enrichment efficiency of the novel method. (b) Percentage recovery was calculated as the percentage of genomes in input DNA that produced DCS reads. CRISPR-DS increases recovery thanks to the initial CRISPR-based target enrichment, which eliminates one round of hybridization capture. (c) After creating DCS reads, the median DCS depth across all targeted regions was calculated for each input amount. The increased recovery enabled by CRISPR-DS translates into 5-10 times more sequencing depth for the same input DNA.

The side-by-side comparison of CRISPR-DS vs standard-DS also demonstrated a substantial increase in recovery using CRISPR-DS. Sequencing recovery, also referred to as yield, is typically measured as the fraction or percentage of sequenced genomes compared to input genomes. Consistent with prior studies[13, 17], standard-DS produced a recovery rate of ~1% across the different inputs, while CRISPR-DS recovery rate ranged between 6 and 12% (Fig. 5b). Notably, 25ng of DNA prepared with CRISPR-DS produced a post-processing depth comparable to 250ng with standard-DS. This indicates that size selection for CRISPR/Cas9 excised fragments not only removes a step from the library preparation but, most importantly, increases the recovery of input DNA enabling deep sequencing with greatly reduced DNA requirements.

### Validation of CRISPR-DS recovery in an independent set of samples, including low quality DNA

We further confirmed the performance of CRISPR-DS in an independent set of 13 DNA samples extracted from bladder tissue (Additional file 2: Table S3). We used 250ng and obtained a median DCS depth of 6,143x, corresponding to a median recovery rate of 7.4% in agreement with the prior experiment. Reproducible performance was demonstrated with technical replicates for two samples (B2 and B4, Additional file 2: Table S3). All samples had >98% reads on-target after consensus making, but the percentage of on-target raw reads ranged from 43% to 98%. We noticed that the low target enrichment corresponded to samples with DNA Integrity Number (DIN) <7. DIN is a measure of genomic DNA quality ranging from 1 (very degraded) to 10 (not degraded) [25]. We reasoned that degraded DNA compromises enrichment by size selection, and the poor yield could be mitigated by removing low molecular weight DNA prior to CRISPR/Cas9 digestion. To test this hypothesis, we used the pulse-field feature of the BluePippin system to select high molecular weight DNA (> 8kb) from two samples with degraded DNA (DINs 6 and 4). This pre-enrichment resulted in successful removal of low molecular weight products and increased on-target raw reads by 2-fold and DCS depth by 5-fold (Additional file 1: Figure S6). These results indicate that enrichment of high molecular weight DNA could be used as a solution for successful CRISPR-DS performance in partially degraded DNA.

### Validation of CRISPR-DS for the detection of low-frequency mutations

To validate the ability of CRISPR-DS to detect low-frequency mutations, we analyzed four peritoneal fluid samples collected during debulking surgery from women with ovarian cancer and previously analyzed for *TP53* mutations using the standard-DS protocol [17]. The tumor mutation was previously identified in the four samples: in one sample at a high frequency (68.5%) and at a very low frequency (around or below 1%) in the remaining 3 samples. CRISPR-DS detected the tumor mutation in all samples at frequencies comparable to what was reported in the original study (Table 2) [17]. In addition to the tumor mutation, standard-DS also revealed the presence of additional exonic *TP53* mutations in these samples which were at an extremely low frequency (<0.1%) in all cases. These mutations are considered “biological background” mutations to distinguish them from the tumor-derived mutations [17]. Standard-DS revealed between 1 to 5 biological background mutations in each of the samples, representing an overall mutation frequency of about ~1x10^−6^. Similarly, CRISPR-DS identified biological background mutations in the 4 samples at a comparable overall mutation frequency (Additional file 1: Figure S7). These results indicate that CRISPR-DS preserves the sequencing accuracy and sensitivity for mutation detection previously described for DS [13, 17].

**Table 2.**
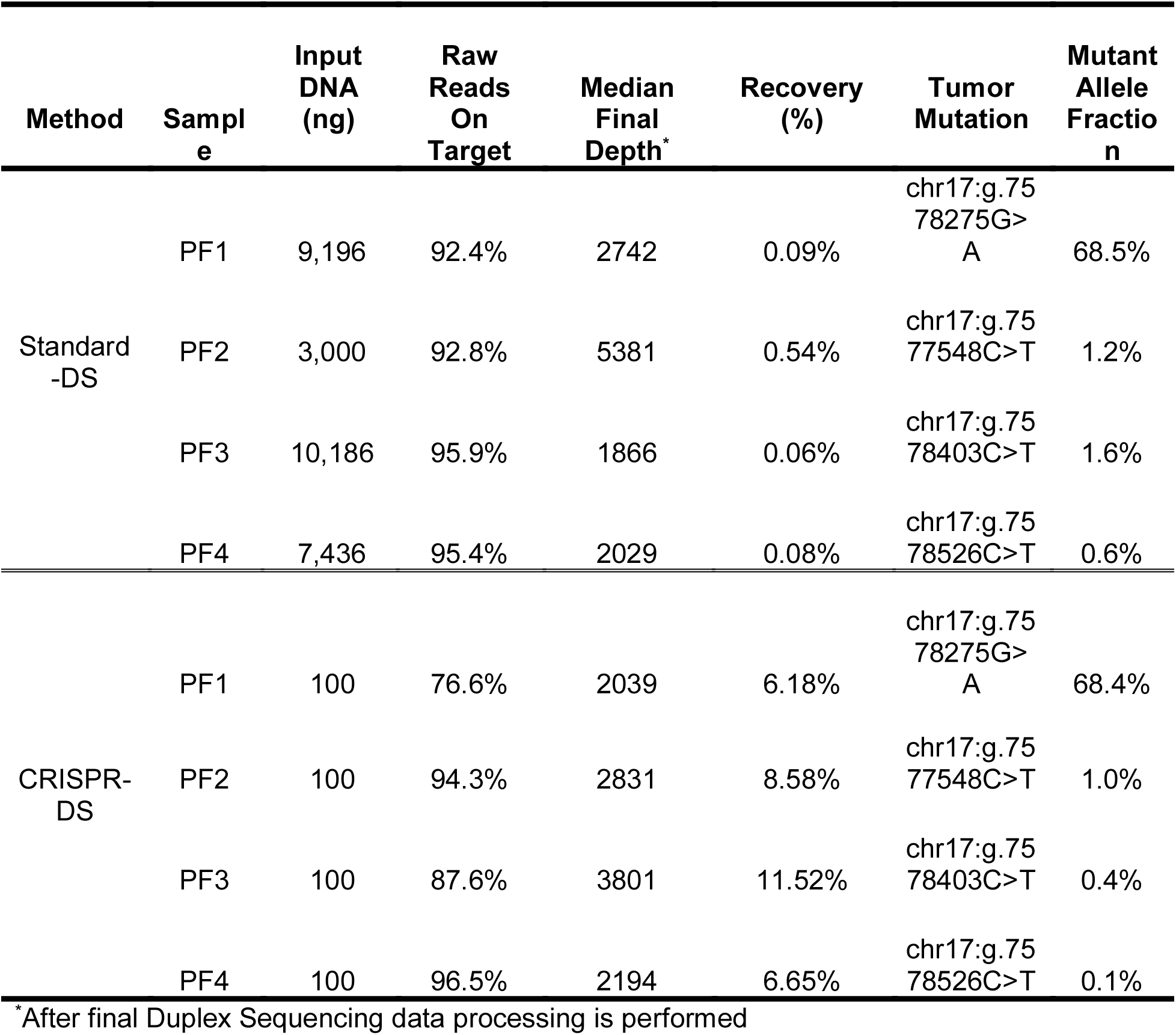
Comparison of Standard-DS vs CRISPR-DS for four different samples with *TP53* mutations.

Table 2 also illustrates a critical advantage of CRISPR-DS compared to standard-DS in terms of translational applicability: the reduced requirement of input DNA as a result of a more efficient library preparation method that enables higher recovery. Standard-DS of these peritoneal fluid samples required between 3-10 μg of DNA to compensate for the ~1% recovery rate of standard-DS and to achieve the high depth necessary to detect low frequency tumor mutations. With CRISPR-DS we only used 100ng of DNA (30-100 fold less than what was used for standard-DS), and we obtained comparable DCS depth to standard-DS (Table 2). Recovery rates ranged between 6 and 12%, as in prior experiments (Fig. 5 and Additional file 2: Table S3). These results represent an efficiency increase of 15x-200x compared to standard-DS with the same DNA. Notably, CRISPR-DS not only preserved sensitivity for mutation detection, increased sequencing recovery, and reduced DNA input, but also shortened the protocol by nearly one day (Additional file 1: Figure S2), making it a more cost effective option for accurate deep sequencing of samples with limited DNA amounts.

## DISCUSSION

While CRISPR-based target enrichment can be applied to any sequencing method that requires hybridization capture of small targets, here we have leveraged its qualities for the optimization of DS, producing a new method called CRISPR-DS. CRISPR-DS merges the increases in efficiency provided by CRISPR-based targeted genome fragmentation with the high accuracy of sequencing provided by double strand molecular barcodes, thus enabling ultra-accurate sequencing of small target regions using minimal DNA inputs. In addition to CRISPR-DS, the CRISPR-based target enrichment approach can be used in combination with other methods for targeted sequencing to improve recovery of small targets and to reduce PCR bias and uneven coverage arising from random fragment sizes.

Targeted sequencing remains a cost effective alternative to whole genome-sequencing, especially when high depth is desired [2]. In multiple applications, such as oncology, the goal is to sequence a small panel of relevant genes with high accuracy in order to find low frequency mutations. While the selected target panel can be amplified by PCR, this method creates uneven coverage and false mutations, thus hybridization capture is typically preferred [2]. Hybridization capture improves coverage uniformity and removes certain artifactual mutations but does not resolve these issues completely. A major disadvantage in hybridization-based sequencing methods is the reliance on sonication for genome fragmentation which generates DNA fragments of random size. We have demonstrated that this size heterogeneity generates two problems that can be solved by replacing sonication with CRISPR-based genome fragmentation. The first problem is PCR bias, which results in the preferential amplification of short DNA fragments. PCR bias leads to wasted reads that contain an excess of PCR copies of the same molecule. While these reads can be removed bioinformatically [26], the amplification advantage of certain molecules can lead to uneven coverage and reduced recovery [27]. In methods that employ molecular barcodes, such as DS, three PCR copies are typically sufficient to generate a consensus sequence [14]. Thus, additional sequencing of PCR copies does not produce additional data and only wastes resources. We have demonstrated that with CRISPR-based fragmentation all fragments amplify similarly. This homogeneous amplification translates into uniform coverage across all targeted regions, a critical feature when the goal is to detect low frequency mutations in selected panel of genes.

The second problem associated with the heterogeneous fragment sizes relates to reduced data yield at the read level. Because sonication allows minimal control over fragment size, a large proportion of fragments are typically too short or too long compared to the optimal length size determined by the number of sequencing cycles. When reads are too short, paired-end reads overlap and the middle region is double-sequenced. Conversely, when reads are too long, the middle part of the DNA fragment, which may contain a variant or region of interest, remains un-sequenced. This inefficient read usage is solved with CRISPR-based target selection because the fragments are tailored to the desired read length.

CRISPR-based target fragmentation also offers two additional advantages. First, homogeneously sized DNA fragments can be visualized to confirm library target enrichment prior to sequencing. In sonication-based hybridization capture, the gel electrophoresis for a target-enriched library looks identical to a library with no target enrichment. This issue can result in the costly waste of a sequencing run where the majority of reads are in off-target regions. We show that the defined fragment lengths created by CRISPR-based digestion produce distinct peaks which are easily visualized and confirm that the sequencing library is target-enriched. A second advantage of Cas9 digestion over sonication is the elimination of sonication-induced sequencing errors [3] and the preservation of double stranded DNA at the ends of fragments. Sonication produces ssDNA at the end of molecules which is susceptible to damage and converted into “pseudo-dsDNA” by end repair. This process has the potential to introduce false variant calls, but it is prevented by CRISPR-DS because Cas9 produces blunt ends which do not require end repair.

In the context of small target sequencing by hybridization-capture, the major advantage introduced by CRISPR-based target enrichment is increased recovery, that is, percentage of input genomes that produce sequencing data. Hybridization capture is notably inefficient, especially for small target regions [19, 20]. As demonstrated with our experiments and in agreement with prior studies, the average recovery rate of DS is ~1% which translates to at least 1 μg of DNA being needed to produce an average depth of ~3,000x. This recovery is improved 10-fold by the addition of CRISPR-based target enrichment and the elimination of one round of capture. We have demonstrated that by simply excising the genomic regions of interest and performing size selection, we can achieve a level of enrichment comparable to a single round of capture. By performing this step prior to library preparation, only one round of hybridization capture is needed, greatly minimizing DNA loss and increasing recovery. Therefore, using CRISPR-based target enrichment prior to DS achieves the same depth with 10 times less DNA.

To take advantage of the accuracy of DS while enabling low DNA inputs, several groups have developed DS-based approaches that combine endogenous and exogenous barcodes. Yet each comes with its own set of compromises. BotSeqS, iDES, and SIP-HAVA-Seq all use DS-based error correction and require little DNA input [12, 28, 29], but the reliance on endogenous barcodes means that depth is limited in order to keep shearing points unique. BiSeqS uses chemical conversions to distinguish one strand from another in combination with molecular barcodes [30] which allows for an increased recovery and high sequencing depth. However, as a consequence of the chemical conversions, it is unable to detect all mutation types. In contrast, CRISPR-DS preserves the sequencing accuracy of DS because it relies exclusively on exogenous double strand molecular barcodes, and the error correction method and analytical algorithms remain identical to standard-DS. We have demonstrated that CRISPR-DS identified very low frequency mutations previously detected by DS, confirming its sensitivity. Remarkably this validation experiment was performed with 10 to 100 times less DNA than the original standard-DS experiment, illustrating a significant improvement in recovery that will enable the use of the extreme sensitivity of DS for mutation detection in samples with low input DNA.

Though CRISPR-DS addresses several needs in targeted NGS, it could still benefit from optimizations. First, improvements could be made to increase the recovery of degraded samples. Currently, in order to perform efficient target enrichment with CRISPR/Cas9 digestion and size selection, degraded samples must be pre-processed to remove low molecular weight fragments. We performed this pre-processing using electrophoretic size selection with the BluePippin system. However, to minimize loss of DNA, high molecular weight DNA could be selected with alternative methods such as micro-column filters. Second, we noticed that the best recovery was achieved with smaller inputs of DNA. Since our goal was to achieve higher depth with smaller amounts of input DNA, this was not problematic. However, further efforts should be directed to improve recovery from larger DNA inputs as well. Lastly, although CRISPR-DS provides an effective solution for small-target region deep sequencing, the method becomes costly for deep sequencing of large genomic regions, an inherent problem of deep sequencing. Nevertheless, fragmentation by CRISPR/Cas9 followed by size selection for fragments as a generic target enrichment technique can easily be scaled to many genomic regions as each region only requires the addition of the appropriate gRNAs for target excision. Thus, CRISPR-DS is ideal for small to moderate size panels (1-100Kb) that require ultra-sensitive mutation detection with minimal DNA inputs.

## CONCLUSION

We have demonstrated that CRISPR/Cas9 fragmentation followed by size selection enables efficient target enrichment, increasing the recovery of hybridization capture and eliminating the need for a second round of capture for small target regions. In addition, it eliminates PCR bias, maximizes the use of sequencing resources, and produces homogeneous coverage. This fragmentation method can be applied to multiple sequencing modalities that suffer from these problems. Here we have applied it to DS in order to produce CRISPR-DS, an efficient, highly accurate sequencing method with significantly reduced input DNA requirements. CRISPR-DS has broad application for the sensitive identification of mutations in situations in which samples are DNA-limited, such as forensics and early cancer detection.

## METHODS

### Samples

The samples analyzed included de-identified human genomic DNA from peripheral blood, bladder with and without cancer, and peritoneal fluid DNA from a prior study [17]. Only peritoneal fluid samples had patient information available, which was necessary to confirm the tumor mutation. The remainder of the study samples were used solely to illustrate technical aspects of the technology, no patient information was available, and interpretation of the mutational status of *TP53* is not reported. Frozen bladder samples were obtained from unfixed or frozen autopsy tissue. DNA was extracted with the QIAamp DNA Mini kit (Qiagen, Inc., Valencia, CA, USA) and it had never been denatured, which is essential to preserve the double-stranded nature of each DNA molecule prior to ligation of DS adapters. DNA was quantified with a Qubit HS dsDNA kit (ThermoFisher Scientific). DNA quality was assessed with Genomic TapeStation (Agilent, Santa Clara, CA) and DNA integrity numbers (DIN) were recorded. Peripheral blood DNA and peritoneal fluid DNA had DIN>7 reflecting good quality DNA with no degradation. Bladder samples, however, were purposely selected to include different levels of DNA degradation. Samples B1 to B13 had DINs between 6.8 and 8.9 and were successfully analyzed by CRISPR-DS (Additional file 2: Table S3). Samples B14 and B16 had DINs of 6 and 4, respectively, and were used to demonstrate pre-enrichment of high molecular weight DNA with the BluePippin system (see below and Additional file 1: Figure S6).

### CRISPR guide design

CRISPR/Cas9 uses a gRNA to identify the site of cleavage. gRNAs are composed of a complex of CRISPR RNA (crRNA), which contains the ~20bp unique sequence responsible for target recognition, and a trans-activating crRNA (tracrRNA), which has a universal sequence [31].To select the best gRNAs to excise *TP53* exons we used the CRISPR MIT design website (http://CRISPR.mit.edu). The selection criteria were: (1) production of fragments of ~500bp covering exons 2-11 of *TP53* and (2) highest MIT website score (Additional file 2: Table S1 and Additional file 3: Data S1). For exon 7, a smaller size fragment was required in order to avoid a proximal poly-T repeat (Additional file 1, Figure S3). We designed a total of 12 gRNA, which excised *TP53* into 7 different fragments (Figure 1a). All gRNA had scores >60. 10 gRNAs were successful with the first chosen sequence and 2 had to be redesigned due to poor cutting. Initially, the quality of the cut was assessed by reviewing the alignment of the final DCS reads with Integrative Genomics Viewer [32]. Successful guides produced a typical coverage pattern with sharp edges in region boundaries and proper DCS depth (Figure 3d). Unsuccessful guides led to a drop in DCS depth and the presence of long reads that spanned beyond the expected cutting point. In order to simplify and speed up the assessment of guides, we designed a synthetic GeneBlock DNA fragment (IDT, Coralville, IA) that included all gRNA sequences interspaced with random DNA sequences (Additional file 3: Data S2). 3ng of GeneBlock DNA were digested with each of the gRNAs using the CRISPR/Cas9 in vitro digestion protocol described below. After digestion, the reactions were analyzed by TapeStation 4200 (Agilent Technologies, Santa Clara, CA, USA) (Additional file 1: Figure S9). The presence of predefined fragment lengths confirms: (1) Proper gRNA assembly (2) The ability of the gRNA to cleave the designed site.

### CRISPR/Cas9 *in vitro* digestion of genomic DNA

The *in vitro* digestion of genomic DNA with *S. pyogenes* Cas9 Nuclease requires the formation of a ribonucleoprotein complex, which both recognizes and cleaves a pre-determined site. This complex is formed with gRNAs (crRNA + tracrRNA) and Cas9. For multiplex cutting, the gRNAs can be complexed by pooling all the crRNAs, then complexing with tracrRNA, or by complexing each crRNA and tracrRNA separately, then pooling. The second option is preferred because it eliminates competition between crRNAs. gRNAs are at risk of quick degradation and repeated cycles of freeze-thawing should be avoided. crRNAs and tracrRNAs (IDT, Coralville, IA) were complexed into gRNAs and then 30nM of gRNAs were incubated with Cas9 nuclease (NEB, Ipswich, MA) at ~30nM, 1x NEB Cas9 reaction buffer, and water in a volume of 23-27 μL at 25°C for 10 min. Then, 10-250ng of DNA was added for a final volume of 30 μL. The reaction was incubated overnight at 37°C and then heat shocked at 70°C for 10 min to inactivate the enzyme.

### Size Selection

Size selection for the predetermined fragment length is critical for target enrichment prior to library preparation. AMPure XP Beads (Beckman Coulter, Brea, CA, USA) were used to remove off-target, un-digested high molecular weight DNA. After heat inactivation, the reaction was combined with a 0.5x ratio of beads, briefly mixed, and then incubated for 3 min to allow the high molecular weight DNA to bind. The beads were then separated from the solution with a magnet and the solution containing the targeted DNA fragment length was transferred into a new tube. This was followed by a standard AMPure 1.8x ratio bead purification eluted into 50 μL of TE Low to exchange the buffer and remove small DNA contaminants.

### A-tailing, and ligation

The fragmented DNA was A-tailed and ligated using the NEBNext Ultra II DNA Library Prep Kit (NEB, Ipswich, MA) according to manufacturer’s protocol. The NEB end-repair and A-tailing (ERAT) reaction was incubated at 20°C for 30 min and 65°C for 30 min. Note that end-repair is not needed for CRISPR-DS because Cas9 produces blunt ends, but the ERAT reaction was used for convenient A-tailing. The NEB ligation mastermix and 2.5μl of DS adapters at 15 μM were added and incubated at 20°C for 15 min according to the manufacturer’s instructions. Instead of relying on in-house manufactured adapters using previously published protocols [13, 14], which tend to exhibit substantial batch-to-batch variability, we used a commercial adapter prototype of the structure shown in Fig. 1c that were synthesized externally through arrangement with TwinStrand Biosciences. The two differences from the previous adapters are: (1) 10bp random double stranded molecular tag instead of 12bp and (2) substitution of the previous 3’ 5bp conserved sequence by a simple 3’-dT overhang to ligate onto the 5’-dA-tailed DNA molecules. Upon ligation, the DNA was cleaned by a 0.8X ratio AMPure Bead purification and eluted into 23 μL of nuclease free water.

### PCR

The ligated DNA was amplified using KAPA Real-Time Amplification kit with fluorescent standards (KAPA Biosystems, Woburn, MA, USA). 50μl reactions were prepared including KAPA HiFi HotStart Real-time PCR Master Mix, 23μl of previously ligated and purified DNA and DS primers MWS13, 5’-AATGATACGGCGACCACCGAG-3’, and MWS20, 5’- GTGACTGGAGTTCAGACGTGTGC-3’ [13, 14] at a final concentration of 2 μM. The reactions were denatured at 98°C for 45 sec and amplified with 6-8 cycles of 98°C for 15 sec, 65°C for 30 sec, and 72°C for 30 sec, followed by final extension at 72°C for 1 min. Samples were amplified until they reached Fluorescent Standard 3, which typically takes 6-8 cycles depending on the amount of DNA input. Reaching Fluorescent Standard 3 produces a sufficient and standardized number of DNA copies into capture across samples and prevents over-amplification. A 0.8X ratio AMPure Bead wash was performed to purify the amplified fragment and eluted into 40μL of nuclease free water.

### Capture and post-capture PCR

*TP53* xGen Lockdown Probes (IDT, Coralville, IA) were used to perform hybridization capture for *TP53* exons as previously reported with minor modifications. From the pre-designed IDT *TP53* Lockdown probes, we selected 21 probes that cover the entire *TP53* coding region (exon 1 and part of exon 11 are not coding) (Additional file 2: Table S2). Each CRISPR/Cas9 excised fragment was covered by at least 2 probes and a maximum of 5 probes (Additional file 3: Data S1). To produce the capture probe pool, each of the probes for a given fragment was pooled in equimolar amounts, producing 7 different pools, one for each fragment. The pools were mixed again in equimolar amounts, except for the pools for exon 7 and exons 8-9, which were represented at 40% and 90% respectively. The decrease of capture probes for those exons was implemented after observing consistent overrepresentation of these exons at sequencing. The final capture pool was diluted to 0.75 pmol/μl. Of note, it is essential to dilute the capture pool in low TE (0.1 mM EDTA) and to aliquot it in small volumes suitable for 2-3 uses. Excessive rounds of freeze-thaw severely impact the efficiency of the protocol. Hybridization capture was performed according to the IDT protocol, except for 3 modifications. First, we used blockers MWS60, 5’-AATGATACGGCGACCACCGAG ATCTACACTCTTTCCCTAC ACGACGCTCTTCCGATCTIIIIIIIIIIIITGA CT-3’ and MSW61, 5’-GTCAIIIIIIIIIIIIAGATCGGAAGAGCACACGTCTGAACTCCAGTCAC-3’, which are specific to DS adapters. Second, we used 75μl of Dynabeads M-270 Streptavidin beads instead of 100μl. Third, the post-capture PCR was performed with the KAPA Hi-Fi HotStart PCR kit (KAPA Biosystems, Woburn, MA, USA) using MWS13 and indexed primer MWS21 at a final concentration of 0.8 μM. The reaction was denatured at 98°C for 45 sec and then amplified for 20 cycles at 98°C for 30 sec, 60°C for 45 sec, and 72°C for 45 sec, followed by extension at 72°C for 60 sec. The PCR product was purified with a 0.8X AMPure Bead wash.

### Sequencing

Samples were quantified using the Qubit dsDNA HS Assay Kit, diluted, and pooled for sequencing. The sample pool was visualized on the Agilent 4200 TapeStation to confirm library quality. The TapeStation electropherogram should show sharp, distinct peaks corresponding to the fragment length of the designed CRISPR/Cas9 cut fragments (Fig. 3b-c). This step can also be performed for each sample individually, prior to pooling, to verify the performance of each individual sample. The final pool was quantified using the KAPA Library Quantification kit (KAPA Biosystems, Woburn, MA, USA). The library was sequenced on the MiSeq Illumina platform using a v3 600 cycle kit (Illumina, San Diego, CA, USA) as specified by the manufacturer. For each sample, we allocated ~7-10% of a lane corresponding to ~2 million reads. Each sequencing run was spiked with approximately 1% PhiX control DNA.

### Standard-DS experiments

Three amounts of DNA (25ng, 100ng, and 250ng) from normal human bladder sample B9 were sequenced with standard-DS with one round and two rounds of capture to provide direct comparison with CRISPR-DS. Standard-DS was performed as previously described [14], with the exception that the KAPA Hyperprep kit (KAPA Biosystems, Woburn, MA, USA) was used for end-repair and ligation and the KAPA Hi-Fi HotStart PCR kit (KAPA Biosystems, Woburn, MA, USA) was used for PCR amplification. Hybridization capture was performed with xGen Lockdown probes that covered *TP53* exons 2-11, the same that were used for CRISPR-DS. Samples were sequenced on ~10% of a HiSeq 2500 Illumina platform to accommodate shorter fragment lengths. Data analysis was perform with the standard-DS analysis pipeline (https://github.com/risqueslab/DuplexSequencingScripts).

### CRISPR-DS target enrichment experiments

Two different experiments were performed to characterize CRISPR-DS target enrichment. The first experiment consisted of comparing one vs. two rounds of capture. Three DNA samples were processed for CRISPR-DS and split in half after one hybridization capture. The first half was indexed and sequenced and the second half was subject to an additional round of capture, as required in the original DS protocol. Then the percentage of raw reads on-target (covering *TP53* exons) was compared for one vs. two captures. The second experiment assessed the percentage of raw reads on-target without performing hybridization capture to determine the enrichment produced exclusively by size selecting CRISPR excised fragments. Fold-enrichment was calculated as the fraction of on-target raw reads divided over the expected fraction of on-target reads given the size of the target region (bases in the target region/total genome bases). Different DNA amounts (from 10ng to 250ng) of three different samples were processed with the protocol described above until first PCR, that is, prior to hybridization capture. Then the PCR product was indexed and sequenced. The percentage of raw reads on-target was calculated and the fold enrichment was estimated considering the size of the targeted region, which is 3,280bp.

### Pre-enrichment for high molecular weight DNA

Selection of high molecular weight DNA improves the performance of degraded DNA in CRISPR-DS. We performed this selection using a BluePippin system (Sage Science, Beverly, MA). Two bladder DNAs with DINs of 6 and 4 were run using a 0.75% gel cassette and high-pass setting to obtain >8kb fragments. Size selection was confirmed by TapeStation (Additional file 1: Figure S6a). Then 250ng of DNA before BluePippin and 250ng of DNA after BluePippin were processed in parallel with CRISPR-DS. The percentage of raw reads on-target as well as average DCS depth was quantified and compared (Additional file 1: Figure S6b.). Alternative methods for size selection such as AMPure beads might be suitable to perform this enrichment.

### Data processing

A custom bioinformatics pipeline was created to automate analysis from raw FASTQ files to text files (Additional file 1: Figure S8). This pipeline includes two major modifications compared to the previously described method for DS analysis [13, 14]: (1) the retention of paired read information and (2) consensus-making performed prior to alignment. Paired-end reads are essential to the analysis of CRISPR-DS data, but are also an important improvement for the analysis of DS in general, as they allow critical quality control of fragment size and removal of potential technical artifacts related to short fragments. In this pipeline, consensus is executed by a custom python and bash scripts. After consensus calling, the resulting processed FASTQ files are aligned to the reference genome of interest, in this case human reference genome v38, using bwa-mem v.0.7.4[33] with default parameters. Mapped reads are re-aligned with GATK Indel-Realigner and low quality bases are clipped from the ends with GATK Clip-Reads (https://software.broadinstitute.org/gatk/). Because of the expected decrease in read quality in the latest cycles of sequencing, we performed a conservative clipping of 30 bases from the 3’ end and another 7 bases from 5’ end were clipped to avoid the occasional extra overhang left by incorrectly synthesized adapters. In addition, overlapping areas of read-pairs, which in our *TP53* design spanned ~80bp, are trimmed back using fgbio ClipOverlappingReads (https://github.com/fulcrumgenomics/fgbio). Software for CRISPR-DS is available at https://github.com/risqueslab/CRISPR-DS.

### Data analysis

Recovery rate (also called fractional genome-equivalent recovery) was calculated as average DCS depth (sequenced genomes) divided by number of input genomes (1ng of human genomic DNA corresponds to ~330 haploid genomes). The number of on-target raw reads was calculated by counting the number of reads within 100bp window on either side of the CRISPR/Cas9 cut sites. Optimal fragment size (Fig. 4b-c and Additional file 1: Figure S4) was calculated as the sequencing read length minus the barcode sequence and minus clipped off bases for poor quality at the ends of reads. For peritoneal fluid samples sequenced with both CRISPR-DS and standard-DS, *TP53* biological background mutation frequency was calculated as the number of *TP53* mutations in *TP53* exons 4 to 10 (excluding the tumor mutation) divided by the total number of nucleotides sequenced in those exons. The 95% confidence intervals were calculated in R using the Clopper-Pearson ‘exact’ method for binomial distributions.

## ABBREVIATIONS

DS: Duplex Sequencing
DCS: Double-stranded consensus sequence
SSCS: Single-stranded consensus sequence
gRNA: Guide RNA
crRNA: CRISPR RNA
tracrRNA: Trans-activating crRNA
NGS: Next-generation Sequencing
ng: Nanogram
bp: Basepair
ssDNA: Single-stranded DNA
dsDNA: Double-stranded DNA
DIN: DNA integrity number

## DECLARATIONS

### Ethics approval and consent to participate

Samples in these studies were obtained from: (1) the University of Washington Gynecologic Oncology Tissue Bank, which collected specimens and clinical information after informed consent under protocol number 27077 approved by the University of Washington Human Subjects Division institutional review board; (2) the University of Washington Genitourinary Cancer Specimen Biorepository and from not previously fixed or frozen autopsy tissue with waiver of consent under protocol number 52389 approved by the Fred Hutchinson Cancer Research Center Human Subjects Division institutional review board.

### Consent for publication

All the samples in the study were de-identified. Consent for publication is included under the informed consent for research described above.

### Availability of data and material

Sequencing data that supports the findings of this study have been deposited in the Sequence Read Archive (BioProject ID: <u>PRJNA412416</u>). Software for CRISPR-DS data analysis is available at https://github.com/risqueslab/CRISPR-DS.

### Competing interests

SRK is a consultant and equity holder for TwinStrand Biosciences Inc. JJS is a founder and equity holder in TwinStrand Biosciences Inc. RAR is the principal investigator on a NIH SBIR R44CA221426 subcontract research agreement with TwinStrand Biosciences Inc.

### Funding

Research reported in this publication was supported by grants from the NIH under award numbers R01CA160674 and R01CA181308 to RAR; Mary Kay Foundation grant 045-15 to RAR. Cooperative Agreement Number W911NF-15-2-0127 from the Department of Defense Army Research Office/Defense Forensic Science Center(DFSC), as well as grant W81XWH-16-1-0579 from the Department of Defense Congressionally Directed Medical Research Program to SRK. The views and conclusions contained in this document are those of the authors and should not be interpreted as representing the official policies, either expressed or implied, of the Army Research Office, DFSC, or the U.S. Government. The U.S. Government is authorized to reproduce and distribute reprints for Government purposes notwithstanding any copyright notation hereon.

### Authors’ contributions

S.R.K. conceived the idea; D.N. S.R.K., and R.A.R. developed the method; D.N. and R.A.R. designed the experiments; D.N., S.L., E.K.S., M.J.H., K.B., K.L.S., B.F.K., R.A.R, and S.R.K. carried out experiments and/or performed data analysis; M.T. provided samples and scientific input; Y.Z., and J.S. contributed to assay development and provided invaluable critical discussion; D.N. S.R.K. and R.A.R. wrote the paper.

## Acknowledgements

We thank Shilpa Kumar for assistance with computational analysis, Emily Kohlbrenner for technical support and helpful discussions, Penny Faires for critical reading and copy editing of the manuscript, and the Genitourinary Cancer Specimen Biorepository for providing access to bladder cases (Director Dr Colm Morrissey, PhD), We thank the University of Washington Gynecologic Oncology Tissue Bank for providing peritoneal fluid DNA and the Brigham and Women’s Hospital/Harvard Cohorts Biorepository for sending archived samples from the Nurses’ Health Study for pilot testing.

## ADDITIONAL FILE 1

### SUPPLEMENTARY FIGURES

Figure S1. Comparison of mutation limit detection by sequencing accuracy

Figure S2. Timeline of library preparation for CRISPR-DS and standard-DS

Figure S3. Homopolymer region produces suboptimal sequencing near *TP53* exon 7

Figure S4. Fraction of reads within 10% of optimal insert size: CRISPR-DS vs standard-DS

Figure S5. Target enrichment for CRISPR-DS with one vs. two captures

Figure S6. Pre-enrichment for high molecular weight DNA with BluePippin

Figure S7. Comparison of *TP53* biological background mutation frequency measured by Standard-DS and CRISPR-DS

Figure S8. Overview of CRISPR-DS data processing

Figure S9. Control CRISPR/Cas9 digestion of *TP53* gRNAs

## ADDITIONAL FILE 2

### SUPPLEMENTARY TABLES

Table S1. crRNA sequences for *TP53* CRISPR/Cas9 digestion

Table S2. *TP53* hybridization capture probes

Table S3. CRISPR-DS sequencing results for 15 samples processed with 250ng input DNA

## ADDITONAL FILE 3

### SUPPLEMENTARY DATA

Data S1. *TP53* sequence with crRNA and capture probes

Data S2. GeneBlock sequence

